# Hydrodynamic synchronization of elastic cilia: How flow confinement determines the characteristics of metachronal waves

**DOI:** 10.1101/2023.10.20.563276

**Authors:** Albert von Kenne, Markus Bär, Thomas Niedermayer

## Abstract

Cilia are hair-like micro-actuators whose cyclic motion is specialized to propel extracellular fluids at low Reynolds numbers. Clusters of these organelles can form synchronized beating patterns, called metachronal waves, which presumably arise from hydrodynamic interactions. We model hydrodynamically interacting cilia by microspheres elastically bound to circular orbits, whose inclinations with respect to the cellular wall model the ciliary power and recovery stroke, resulting in an anisotropy of the viscous flow. We derive a coupled phase oscillator description by reducing the microsphere dynamics to the slow time scale of synchronization and determine analytical metachronal wave solutions and their stability in a periodic chain setting. In this framework, a simple intuition for the hydrodynamic coupling between phase oscillators is established by relating the geometry of near-wall flow to the directionality of the hydrodynamic coupling functions. This intuition naturally explains the properties of the linear stability of metachronal waves. The flow confinement at the wall stabilizes metachronal waves with long wavelengths propagating in the direction of the power stroke and, moreover, metachronal waves with short wave lengths propagating perpendicularly to the power stroke. Performing simulations of phase oscillator chains with periodic boundary conditions, we indeed find that both wave types emerge with a variety of linearly stable wave numbers. In open chains of phase oscillators, the dynamics of metachronal waves is fundamentally different. Here, the elasticity of the model cilia controls the wave direction and selects a particular wave number: At large elasticity, waves traveling in the direction of the power stroke are stable, whereas at smaller elasticity waves in the opposite direction are stable. For intermediate elasticity both wave directions coexist. In this regime, waves propagating towards both ends of the chain form, but only one wave direction prevails, depending on the elasticity and initial conditions.

## I. INTRODUCTION

A generic way of living matter to achieve fluid propulsion is the cyclic motion of cilia and eukaryotic flagella [1]. These organelles are hair-like cell extensions composed of a circular array of microtubule-doublets called an axoneme [2]. The axonemal structure is subject to an internal biochemistry that drives it into a cyclic motion [3] characterized by an asymmetry known as the power and recovery stroke [4] – power indicating the production of thrust, recovery indicating the return to the initial state. This asymmetry is crucial because microscopic flow characteristics require non-reciprocal motion to generate propulsion [5]. The flows generated by cilia and flagella provide essential functions, e. g., single cell locomotion [6], feeding of marine invertebrates [7], mucus clearance in the respiratory tract [8], circulation of cerebrospinal fluid [9] or breaking the symmetry of organ arrangement in developing embryos [10]. Cilia and eukaryotic flagella are structurally identical [1], thus, we use the terms cilia and flagella interchangeably.

In order to transport fluid efficiently, arrays of cilia beat in a strikingly coordinated fashion. Often a constant phase shift between the beating of neighboring cilia is formed and thus traveling waves, called metachronal waves, arise [11]. While these waves are ubiquitous in biology, covering scales from microns to centimeters [12, 13], the characteristics of metachronal coordination appear to be specific: A symplectic wave (for which the wave direction is identical to the direction of the power stroke) has been observed in the green algae Volvox carteri [14, 15]. Experiments with reef coral larvae show laeoplectic metachronal waves (for which the wave propagates to the left of the power stroke) [16]. The metachronal waves in the unicellular ciliate Paramecium propagate to the right of the power stroke (dexioplectic) [17, 18]. In ciliated epithelia studied in vitro, symplectic, laeoplectic and oblique waves occur depending on mucus conditions [19].

The physics governing the interactions between cilia is a widely-discussed topic. Plausible coupling contributions are mechanical connections [20], steric interactions [21, 22] and, notably, hydrodynamic forces mediated by the extracellular fluid [23]. Empirical evidence on this issue supports different effects depending on the specifics of the biological system: Coupling two isolated flagella extracted from the somatic cells of flagellated algae through a fluid medium shows that hydrodynamic interactions alone can lead to in-phase motion, anti-phase motion and non-trivial phase locking [24, 25]. On the other hand, subjecting the flagella of Chlamydomonas reinhardtii to cyclic external flows and studying the phase locking properties suggests that the magnitude of hydrodynamic forces in Chlamydomonas is insufficient to explain synchronization, pointing to mechanical couplings [26]. A similar analysis, however, indicates that hydrodynamic interactions could be the dominant coupling between cilia in mammalian brain tissue [27].

Detailed numerical simulations of hydrodynamic interactions between beating cilia generally show the emergence of synchronization patterns [28–30] and suggest an energetic advantage of metachronal waves [31, 32]. However, as the complexity of these models makes it rather difficult to understand the underlying effects, reduced approaches that model the cilia as self-sustained phase oscillators have been successful to predict synchronization and metachronal waves [33–36]. It turns out that hydrodynamic synchronization requires breaking the time-reversal symmetry of the microscopic flow [37], which can result from elastic waveform compliance [34, 38], substrate-modified viscous drag [33], phase-dependent intrinsic forcing [35], hydrodynamic memory [39], cell body motion [40], or combinations of these mechanisms [41]. In fact, the dominant physical mechanism is still under debate [42, 43].

To investigate the role of hydrodynamic coupling between cilia, the characteristics of metachronal waves are often studied. With the exception of a paper investigating the effect of a spherical substrate [44], recent efforts to construct phase oscillator models of metachronal coordination typically ignore elastic compliance and focus on the details of beating [45–49]: In a carpet of cilia, different metachronal waves are stable, depending on the characteristic forcing and the geometry of the carpet [46], a net flow can spontaneously be generated by the collective motion induced by variable driving forces [49] and the specific shape of the beating cycle can select a dominant metachronal wave [47, 48]. However, the symplectic metachronal wave observed in Volvox carteri has been successfully predicted with an elastic compliance model, in which the flagella are modeled as coupled phase-amplitude oscillators [14, 15, 50].

In this paper, we try to advance the elastic compliance framework in the spirit of the simple phase oscillator model [34]. To this end we consider the anisotropy of viscous flow near a no-slip wall, that results from motion perpendicular to the wall. Following previous work [14], we model cilia as microspheres elastically bound to circular orbits, whose inclinations with respect to the cell wall model the power stroke of cilia. We reduce this model to coupled phase oscillators in Sec. II A and develop a simple geometric intuition for the hydrodynamic coupling between the phase oscillators in Sec. II B. Based on this intuition, the linear stability characteristics of metachronal waves, which we calculate in Sec. III, can be understood as a natural result of flow confinement at the cell wall. With numerical simulations we address the emergence of metachronal waves in periodic (Sec. IV) as well as in open (Sec. V) phase oscillator chains. Our results show that phase oscillators with elastic orbits, coupled by confined flow fields, robustly coordinate metachronal waves, regardless of the boundary and initial conditions.

Furthermore, we show for the first time that bistability of metachronal wave directions can lead to hysteresis and that the tuning of the elasticity of the orbits can be used to switch between the wave directions. Our model exhibits analytic solutions, allows interpretations in terms of simple intuitions, is highly numerically efficient and satisfies phenomenological key requirements, i. e., the generation of directed fluid transport and the robust formation of metachronal waves. Although we introduce the model in the most simple case of phase oscillator chains, it can be easily generalized to oscillator carpets and, in principle, can be used to address a variety of biologically relevant situations. We discuss the interrelations between our results and the literature on synchronization of cilia in Sec. VI.

## II. MODEL

We abstract the cyclic bending wave dynamics of a cilium into the motion of a microsphere, which represents the application of a point force [51]. This approach allows the analytical calculation of the cyclic flow fields in the extracellular fluid, through which the cilia are coupled. Although this approximation appears rather crude, it can accurately capture the average flow produced by flagellated microorganisms [52]. The elastic properties of the cilia are modeled by a harmonic potential centered on a reference radius of circular motion, allowing only modulations in the radius of motion [34]. This model of cilia results in a dynamical description parameterized by the radial amplitude *R*(*t*) and the phase *ϕ*(*t*). A limit cycle reconstruction based on the analysis of the shapes of the beating flagella of Chlamydomonas and their exposure to controlled flow perturbations motivates this type of model [53].

The fluid flow around the cilia is dominated by the effects of viscosity, as indicated by the low Reynolds number Re = *L*^2^*/νT* ≪ 1, where *L* is the typical cilia length, *T* is the typical period, and *ν* is the kinematic viscosity [54]. Thus, the inertial terms in the Navier-Stokes equations will be neglected relative to the viscous terms. Therefore, the flow field is determined by the stationary Stokes equations

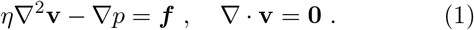

Here, **v**(**r**) and *p*(**r**) are the velocity and pressure fields, *η* is the dynamic viscosity and ***f*** is a body force. Since Eq. (1) lacks explicit time dependence the instantaneous configuration of the cilia completely determines the flow field. For the flow induced by the motion of a small sphere, at distances much larger than its size, the body force can be approximated by a point force **f** *δ*(**r** − **r**_**f**_). The flow velocity is linearly related to the force exerted as

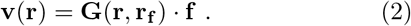

The Greens function **G** of Eq. (1) subject to a no-slip condition at a planar wall is known as the Blake tensor [55]. We use the Blake tensor to describe the hydrodynamic interaction.

The model outlined above can be reduced to coupled phase oscillators by exploiting the separation of time scales that occurs for weak hydrodynamic interactions, namely between fast oscillations and slow synchronization on the one hand, and between fast elastic relaxation and slow synchronization on the other hand [34, 43, 56]. To extract the slow dynamics of synchronization, the dynamic equations of phase and radial amplitude are cycle-averaged. At this level, the radial amplitude varies slowly on the synchronization time scale *t*_s_. Because the time scale of elastic recoil *t*_e_ is small compared to *t*_s_, the radial changes are approximately instantaneous with respect to the synchronization dynamics, reducing the system to coupled phase oscillators that include the effect of orbit compliance as a contribution to the instantaneous phase velocity.

Previous work [34] considered such a phase oscillator model, for orbits oriented parallel to the no-slip boundary. No directed net flow is generated in this configuration. Furthermore, in simulations metachronal waves emerge in periodic chains of phase oscillators, but not in open chains.

Brumley et al. [14, 15] developed a model based on similar elastically bound mechanics and overdamped hydrodynamics. This model retains the dynamics of the radial amplitude. Moreover, a number of additional effects are included in this model, e. g., a non-circular limit cycle, 3D orbit perturbations, and, in particular, the confined flow fields associated with inclined orbits near a no-slip wall. This more general phase-amplitude model generates a net flow and exhibits emergence of metachronal waves independent of the boundary conditions.

We generalize the phase oscillator model [34] by considering inclined orbits near a no-slip wall. The inclination of the orbit breaks the symmetry of circular motion. In the part of the inclined orbit far from the no-slip wall, the microsphere generates a higher fluid velocity, compared to the motion close to the wall. As a result, a directed net flow is generated. In this way, the power stroke of cilia is modeled. The interplay between the flow anisotropy due to the power stroke and the compliance of the orbits is sufficient for robust metachronal wave formation, as we will demonstrate.

While cilia typically occur in two-dimensional arrays, we confine the treatment of the model below to onedimensional chains and investigate longitudinally spreading metachronal waves that propagate in or against the direction of the power stroke, as well as transversally spreading metachronal waves that move orthogonal to the power stroke. The former class is known as symplectic or antiplectic waves, respectively, whereas the literature refers to the latter wave type as laeoplectic or dexioplectic.

### A. Dynamic equations

We model a chain of cilia by a collection of *N* microspheres with radius *a* that are elastically bound to a circular orbit (Fig. 1). The position of the *i*th sphere (*i* ∈ [0, …, *N* − 1]) is written as

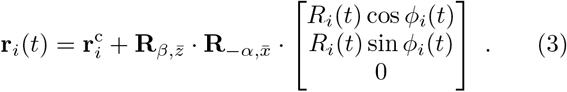

**FIG. 1.**
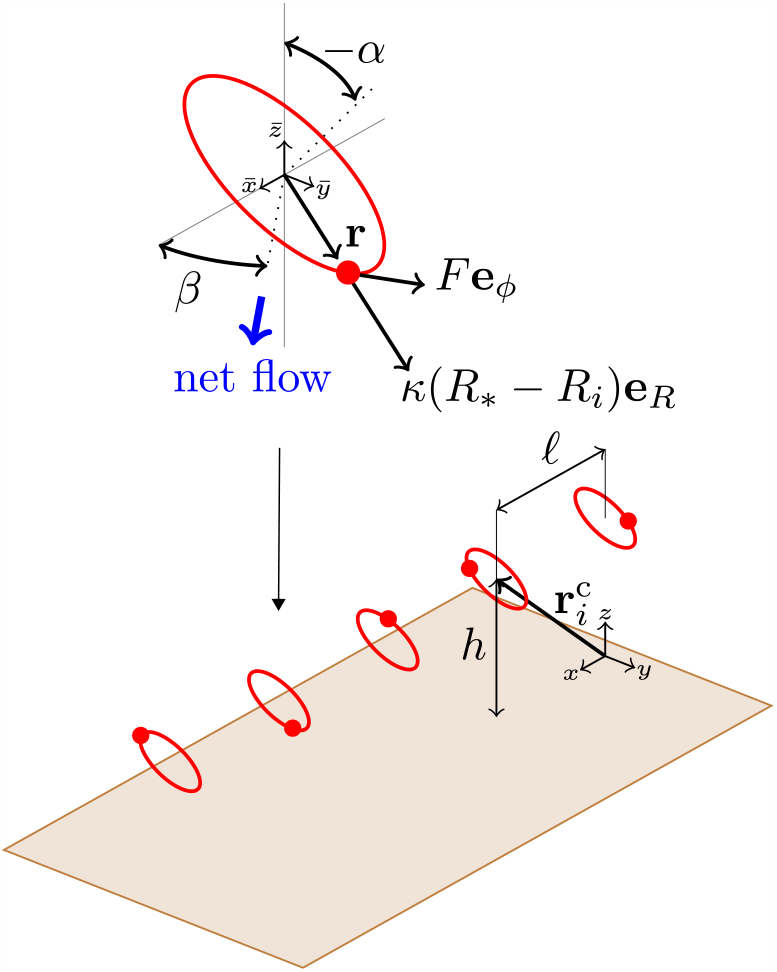
Oscillator model of a cilia chain. A constant tangential force *F* **e**_*ϕ*_ drives microspheres that are elastically bound to a circular orbit with reference radius *R*_*∗*_, by a harmonic force *κ*(*R*_*i*_ *− R*_*∗*_)**e**_*R*_. The centers 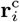 of the orbits are uniformly spaced by a distance . ℓ along the *x*-axis, at a distance *h* above the no-slip wall. The orbits are locally rotated around the construction axis of the oscillator chain, by the rotation matrix 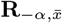. This creates an inclination of the orbits about the substrate by an angle *α*. The orbits are oriented at an angle *β* to the construction axis by a subsequent rotation around the local vertical axis through the rotation matrix 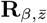. As a consequence of the inclination of the orbits about the substrate a net flow is generated, whose direction is controlled by the orientation of the orbits with respect to the chain axis. The direction of net flow is interpreted as power stroke direction.

Here, 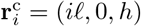 are the centers of the orbits, where *h* denotes the distance to the no-slip wall and *ℓ* is the separation between the center points in the substrate plane.

The rotation matrices 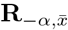 and 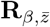 are chosen to create an inclination of the orbits about the wall by an angle *α* and to put the net flow direction at an angle *β* with the construction axis of the oscillator chain, respectively (Fig. 1).

The motility of the cilia is modeled by a constant tangential driving force

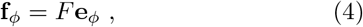

with magnitude *F* . A harmonic restoring force

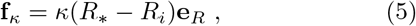

with spring constant *κ* models the elasticity of the cilia and bounds the radial motion to a reference radius *R*_*∗*_. The fluid exerts a drag force on the spheres, which is given by Stokes law

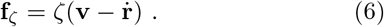

Here, *ζ* = 6*πηa* and **v** is the background velocity field.

In the overdamped Stokes flow, the balance between active, elastic and viscous forces, **f**_*ϕ*_ + **f**_*κ*_ + **f**_*ζ*_ = **0**, leads to the dynamic equations

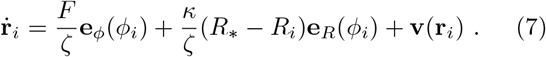

To calculate the fluid velocity at the *i*th sphere **v**(**r**_*i*_), we assume that the size of the sphere is much smaller than the radius of motion *a* ≪ *R*_*∗*_ and that the radius of motion is much smaller than the typical distances *R*_*∗*_ ≪ *ℓ* and *R*_*∗*_ ≪ *h*. Then, the flow velocity can be calculated according to

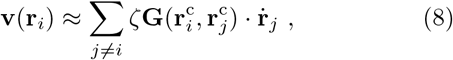

where **G** is the Blake tensor (in App. A we discuss the Blake tensor in detail).

With the flow field Eq. (8) we can write the dynamic equations Eq. (7) as

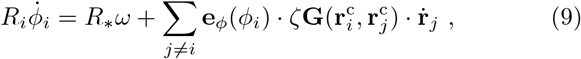

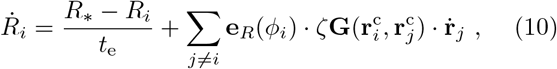

where *ω* = 2*π/T* = *F/ζR*_*∗*_ is the intrinsic frequency and *t*_e_ = *ζ/κ* is the time scale of elastic relaxation.

We assume that the synchronization time *t*_s_ is considerably larger then the period *T* ≪ *t*_s_, as well as the elastic relaxation time *t*_e_ ≪ *t*_s_. In App. A, we show that under these assumptions the dynamics Eqs. (9)-(10) can be reduced to a system of coupled phase oscillators, which reflects the hydrodynamic coupling at the slow time scale *t*_s_. To the leading order in the hydrodynamic interaction we obtain

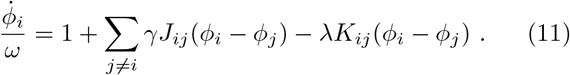

Here, the hydrodynamic interactions between rigid orbits are captured in *γJ*_*ij*_, where *γ* = 3*a/*4ℓ ≪ 1, and the effect of orbit compliance is contained in *λK*_*ij*_, where *λ* = *γt*_e_*ω* ≪ 1.

The parameter *λ/γ* = *t*_e_*ω* indicates the flexibility of the orbits. For *t*_e_*ω* ≪ 1 the orbits are stiff; for *t*_e_*ω ≳* 1 the orbits are soft.

In the following we analyze Eq. (11) with periodic as well as open-ended boundary conditions. For periodic boundary conditions we implement a cutoff radius of half the system size. An open-ended boundary is to say that the system ends at the first and last oscillator, respectively. In this case we impose no cutoff.

### B. Hydrodynamic coupling functions

In App. A, we calculate the hydrodynamic coupling functions at the synchronization time scale. The cycle-averaged hydrodynamic coupling between rigid orbits can be written as

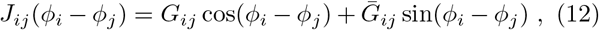

where the geometry factors

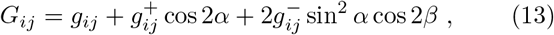

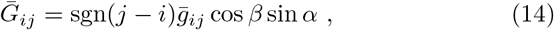

depend on the normalized distance functions

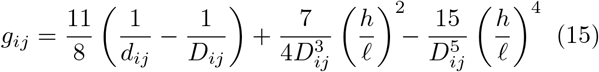

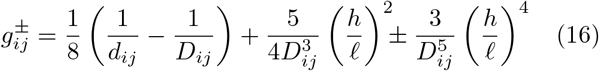

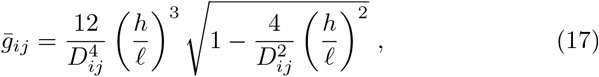

with *d*_*ij*_ = |*i* − *j*| and 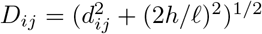.

Notice the symmetry of the geometry factors with respect to the exchange of the phase oscillators: *G*_*ij*_ = *G*_*ji*_ and 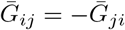.

The cycle-averaged coupling function induced by orbit compliance is given by

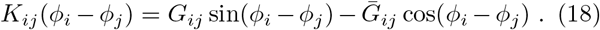

The symmetry of the coupling functions with respect to the exchange of the phase oscillators governs their phenomenological significance. The hydrodynamic interactions between rigid orbits are even:

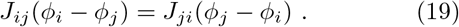

The effect of orbit compliance generates an odd coupling function:

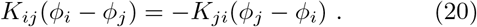

The odd coupling leads to the adaption of the relative phases, which induces synchronization and the formation of metachronal waves. The even coupling causes a frequency gain, representing the cooperative reduction of hydrodynamic drag in the synchronized state.

We can develop an intuition for the characteristics of the hydrodynamic coupling by considering the geometry of flow near the no-slip wall. It is instructive to envisage two particular arrangements of the chain of phase oscillators:

(i) Fig. 2 illustrates the hydrodynamic interaction for an arrangement of phase oscillators whose power stroke direction coincides with the orientation of the chain axis. The flow fields relevant for the coupling lie entirely in the *xz*-plane. Due to the linearity of the Stokes equations it is sufficient to consider the flows generated by a parallel (**f** ^‖^) and a perpendicular force (**f** ^⊥^), as indicated by the red arrows in the sketch of Fig. 2. The flow patterns of Fig. 2(a)-(b) reveal an important anisotropy of flow between the upstream and downstream direction: The flow generated by the parallel force pushes the downstream neighbors away from the substrate, but the upstream neighbors toward the substrate. Similarly the downstream neighbors are pushed in the downstream direction by the flow due to the perpendicular force, but the neighbors upstream are pushed in the upstream direction. This flow anisotropy is reflected in the cycle-averaged coupling by the anti-symmetric geometry factor 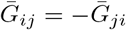. As shown in Fig. 2(c) only the horizontal velocity for the perpendicular force, and the vertical velocity for the parallel force are anisotropic. The other velocity components treat the upstream and downstream neighbors symmetrically, corresponding to a symmetric geometry factor *G*_*ij*_ = *G*_*ji*_. Due to the flow anisotropy, the hydrodynamic interactions between the phase oscillators are balanced at an offset: Two oscillators moving in unison exert partially opposite hydrodynamic forces on each other. Thus, one of the oscillators is pushed to a higher orbit, while the other is pushed to a lower orbit. The resulting opposite changes in the phase velocities drive the hydrodynamically stable configuration away from in-phase motion. The symmetric and asymmetric coupling is balanced at the phase shift

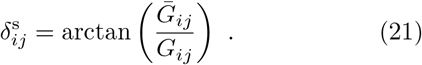

**FIG. 2.**
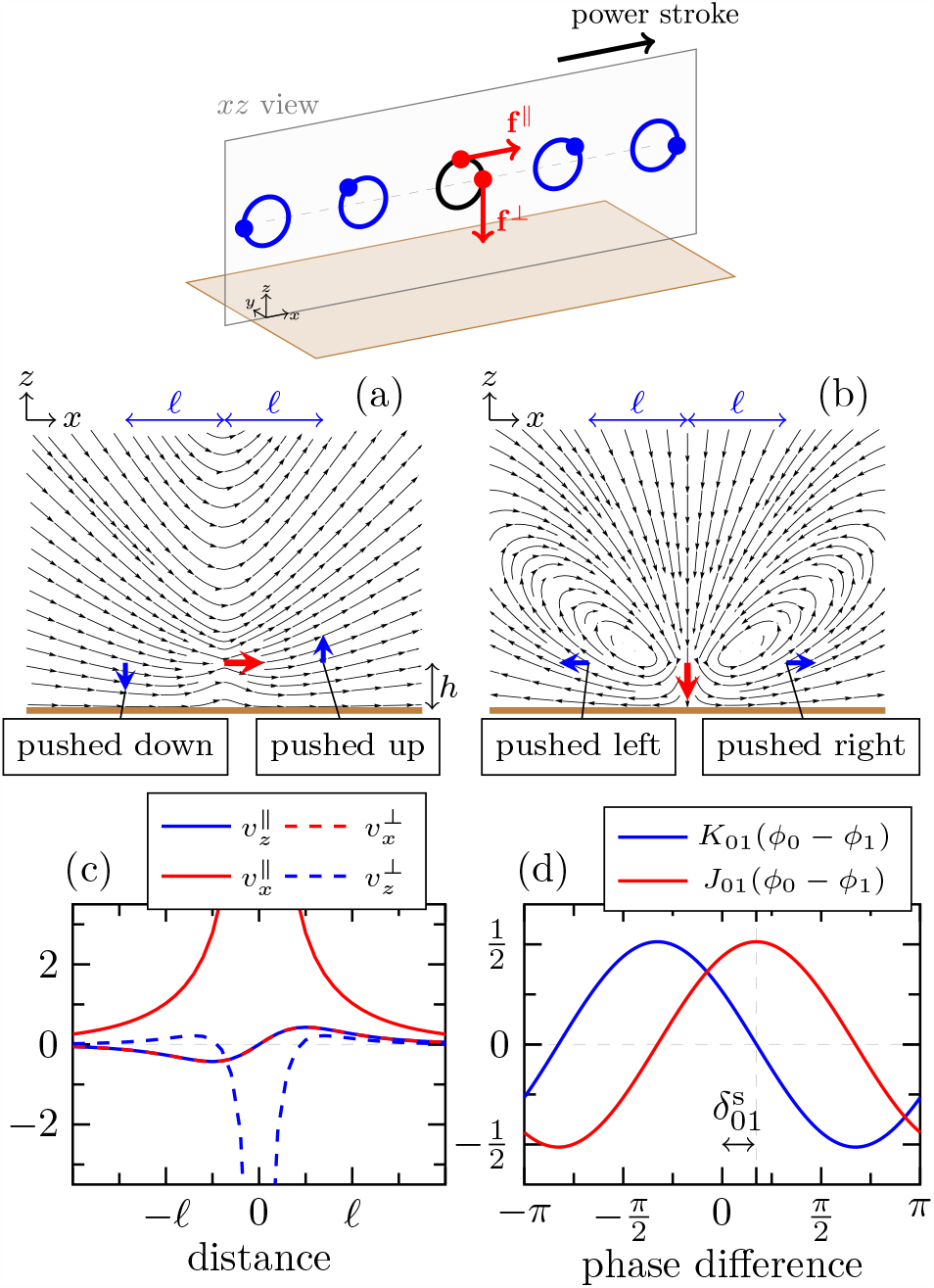
Hydrodynamics for an arrangement of phase oscillators with the power stroke parallel to the chain axis, corresponding to *α* = *π/*2 and *β* = 0. (a) Flow field **v**^‖^ (**r**), according to Eq. (2), caused by a parallel force **f** ^‖^. (b) Flow field **v**^⊥^(**r**) induced by a perpendicular force **f** ^⊥^. (c) Normalized components of the flow velocity generated by the parallel and perpendicular forces, respectively, as a function of distance. (d) Cycle-averaged next-neighbor coupling functions according to Eqs. (12)-(18), with *h/*.ℓ = 1*/*2. The flow anisotropy with respect to upstream and downstream interactions in (a)(b) phase delays the cycle-averaged hydrodynamic coupling by a phase shift 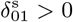.

The corresponding phase shifted cycle-averaged hydrodynamic coupling functions are shown in Fig. 2(d).

(ii) Fig. 3 shows the hydrodynamic interaction for an arrangement of phase oscillators whose power stroke direction is perpendicular to the chain axis. The relevant flow fields lie in the *xz*-plane for vertical perturbations of the phase oscillator motion, and in the *yz*-plane for horizontal perturbations. The power stroke direction is orthogonal to the chain axis, i. e., the hydrodynamic interactions are entirely symmetrical and 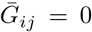. The coupling through the vertical flow velocity due to the vortex shown in Fig. 3(a), however, is qualitatively different for interactions at close and far distances. Close to the forcing (in front of the vortex), the vertical velocity is aligned with the forcing direction. But far from the forcing (beyond the vortex), it is opposite to the forcing direction. The horizontal flow shown in Fig. 3(b) is always in the direction of the forcing. If the vertical flow velocity dominates the horizontal velocity [Fig. 3(c)], the reversal of the vertical flow with respect to the phase oscillator motion changes the hydrodynamically stable configuration between neighbors from in-phase to anti-phase synchronization. In this case, the average coupling functions, depicted in Fig. 3(d), exhibit a phase shift 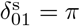.

**FIG. 3.**
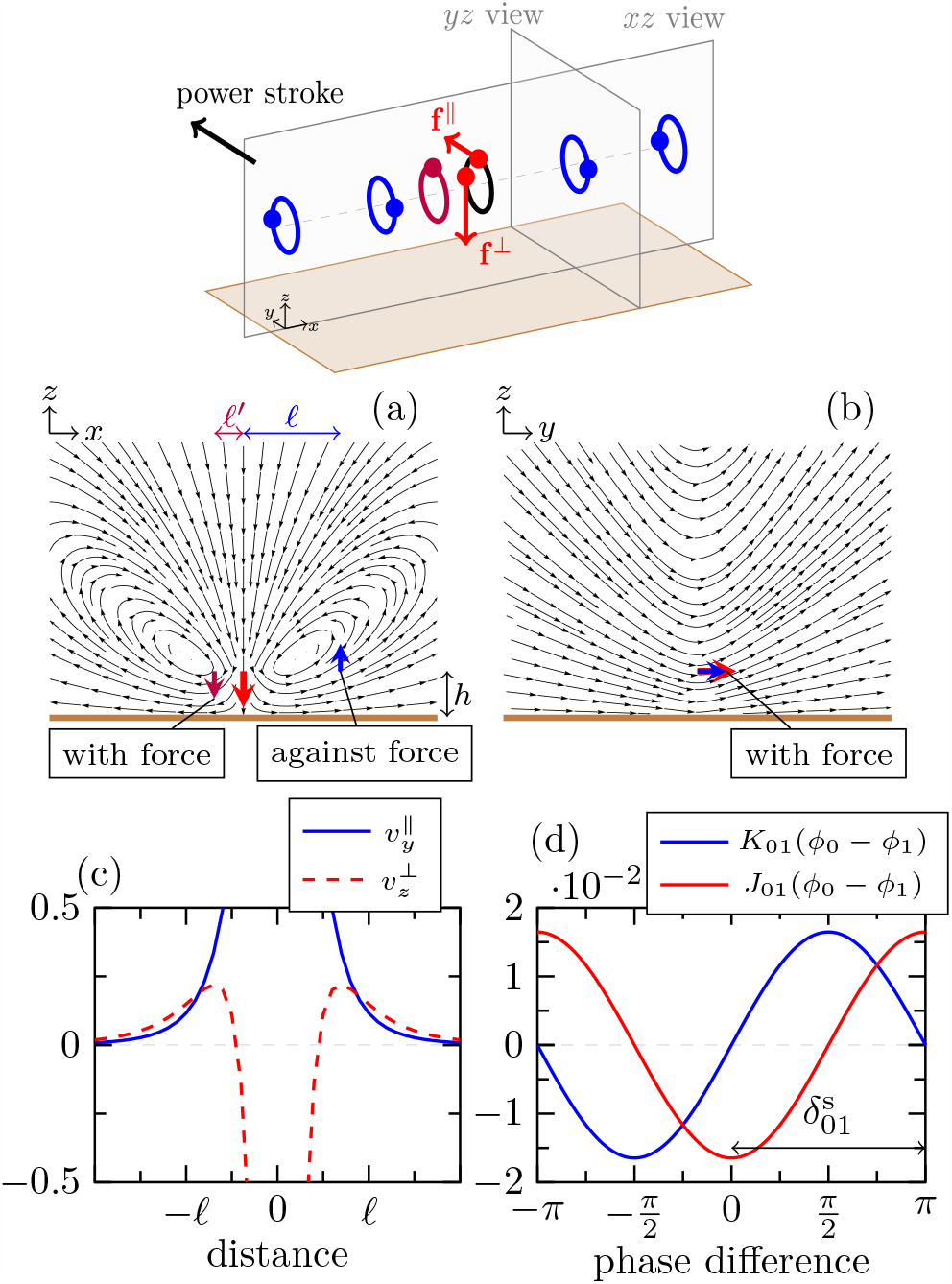
Hydrodynamics for an arrangement of phase oscillators with the power stroke perpendicular to the chain axis, corresponding to *α* = *π/*2 and *β* = *π/*2. (a) Flow field **v**^⊥^(**r**), according to Eq. (2), caused by a perpendicular force. (b) Flow field **v**^‖^ (**r**) caused by a parallel force. (c) Normalized components of the flow velocity generated by the parallel and perpendicular forces, respectively, as a function of distance. (d) Cycle-averaged next-neighbor coupling functions according to Eqs. (12)-(18), with *h/*.ℓ = 1*/*2. The reversal of the vertical flow velocity relative to the direction of the forcing in (a) leads to the sign reversal of the cycle-averaged hydro-dynamic coupling functions, corresponding to a phase shift 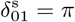.

## III. LINEAR STABILITY OF METACHRONAL WAVES IN PERIODIC CHAINS

We define metachronal waves as frequency-locked solutions of Eq. (11), in which phase shifts *ϕ*_*i*_ − *ϕ*_*i*+1_ remain constant over time but may vary moderately along the chain of phase oscillators.

In this section, we consider chains of *N* phase oscillators with periodic boundary conditions and calculate the linear stability of uniform metachronal waves, where *ϕ*_*i*_ − *ϕ*_*i*+1_ = ℓ*k*, with the wave number *k*. Since the number of phase oscillators is finite, the phase shifts are quantized as |ℓ*k*| = 2*πK/N*, where *K* is an integer. The relationship between metachronal waves in the oscillator phase and waves in continuous media is shown in Fig. 4. Two particular arrangements of the phase oscillators model the basic metachronal waves discussed in biology texts [11]. In an arrangement an as in Fig. 2, the power stroke is directed parallel to the axis of the oscillator chain. Thus, the metachronal waves propagate either with or against the power stroke direction. If the wave direction aligns with the direction of the power stroke, the metachronal waves are called symplectic. For the opposite direction, the metachronal waves are called antiplectic. In the definitions we use, symplectic waves have positive wave numbers ℓ*k >* 0 and antiplectic waves have negative wave numbers ℓ*k <* 0. In an arrangement as in Fig. 3, the direction of the power stroke is perpendicular to the chain axis. Consequently, the wave direction is perpendicular to the power stroke direction. Here, metachronal waves are called dexioplectic if they travel to the right of the power stroke (ℓ*k >* 0) and laeoplectic if they travel to the left (ℓ*k <* 0).

**FIG. 4.**
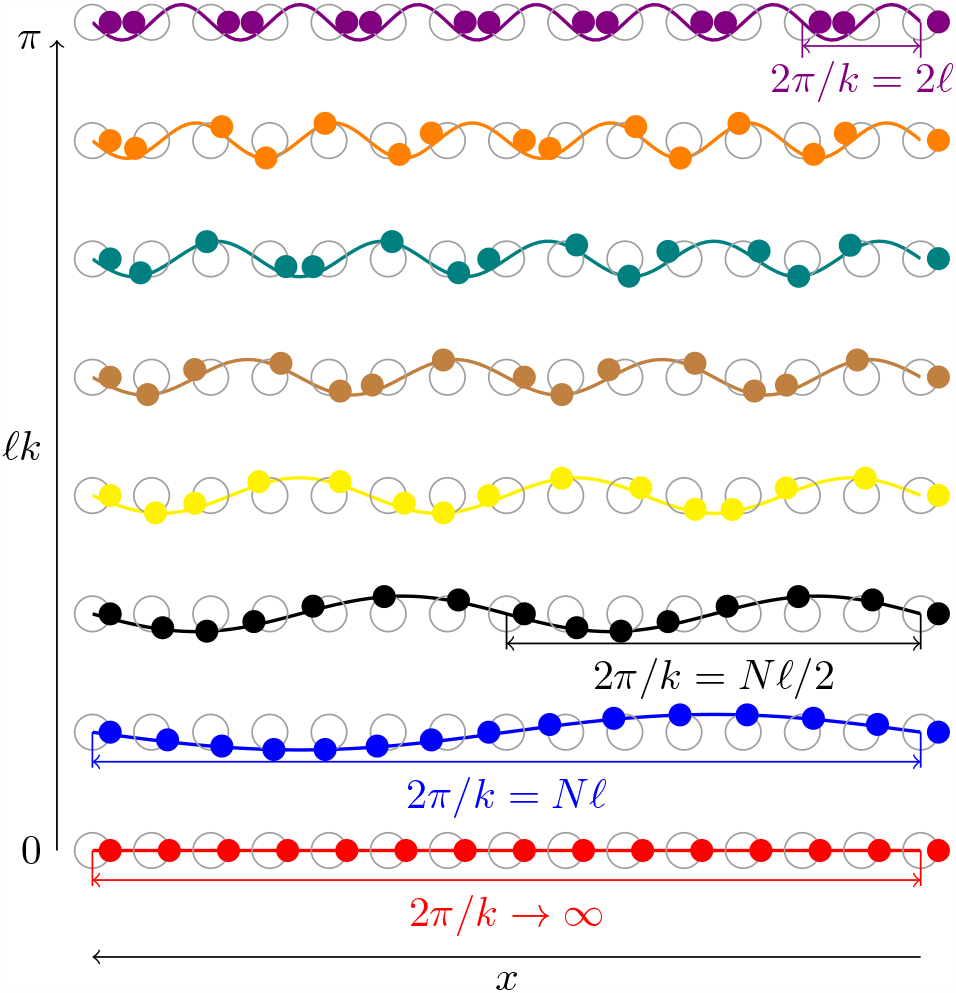
Snapshot of metachronal waves showing the relationship between the collective motion of phase oscillators with phase shifts *ϕ*_*i*_ *− ϕ*_*i*+1_ = . ℓ*k* and a continuous wave sin(*kx*). The global in-phase synchronization corresponds to an infinite wavelength 2*π/k* → ∞ and the global anti-phase synchronization corresponds to a minimum wavelength 2*π/k* = 2. ℓ. We categorize the wave numbers |.ℓ*k*| *> π/*2 as short wavelengths and the wave numbers |.ℓ*k*| *< π/*2 as long wavelengths.

In order to study metachronal waves we work in a corotating frame of reference. The phase in this frame satisfies *ψ*(*t*) = *ϕ*(*t*) − [*ω* + Ω(*k*)]*t*, where the frequency gain of metachronal waves, with wave number *k*, is given by

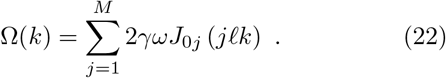

Here, *M* = (*N* − 1)*/*2 for odd *N* and *M* = *N/*2 − 1 for even *N* . The frequency gain is obtained from Eq. (11), with phase shifts *ϕ*_*i*_ −*ϕ*_*i*+1_ = ℓ*k*, using Eqs. (19)-(20) and the periodic boundary conditions.

The dynamic equations of the co-rotating phases are written as

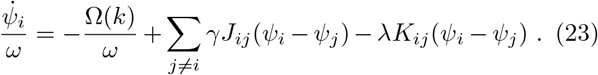

For phases *ψ*_*i*_ − *ψ*_*i*+1_ = ℓ*k* the sum in Eq. (23) gives Ω*/ω*. Thus, metachronal waves are stationary solutions of Eq. (23).

We obtain the dynamics of a small perturbation of metachronal waves **Δ*ψ***(*t*) = [Δ*ψ*_0_(*t*), …, Δ*ψ*_*N−*1_(*t*)], by linearization of Eq. (23) as

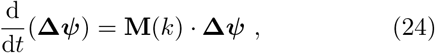

where the components of the Jacobi matrix **M**(*k*) are given by

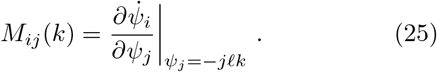

The perturbation exhibits exponential growth in the eigendirections of **M**. The rate of growth in the direction of the *n*the eigenvector is given by the real part of the corresponding eigenvalue Λ_*n*_(*k*). In App. B, we exploit the circulant structure of **M** to calculate the linear growth rates as

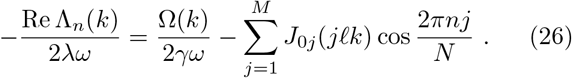

The sign of the linear growth rates determines whether the perturbation grows or decays with time. If each Re Λ_*n*_(*k*) *<* 0 the perturbation exponentially decays in all eigendirections and the metachronal wave with the wave number *k* is linearly stable. The 0th eigenvalue of the circulant matrix **M** is real and vanishes for all wave numbers Λ_0_(*k*) = 0. The eigenvector corresponding to the neutral eigenvalue Λ_0_ is unity for all Δ*ψ*_*i*_, reflecting the invariance against global phase shifts of the underlying model Eq. (23) [34]. Note also, by Eq. (22) and Eq. (26), that the choice of the parameters *λ >* 0, *γ >* 0 and *ω >* 0 does not effect the sign of the linear growth rate.

In order to discuss the linear stability of metachronal waves we first consider next-neighbor coupling. In this case we truncate the sum over pair interactions at *j* = 1, and obtain the linear growth rates as

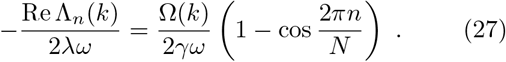

Discarding Λ_0_, the sign of each growth rate is governed by the frequency gain. Thus, if the wave number *k* corresponds to an increased beat frequency, the perturbation decays and the metachronal wave is linearly stable. For orbits oriented in the plane of the substrate the metachronal waves with increased frequency have wave numbers |ℓ*k*| ≤ *π/*2 [34]. If the orbits are inclined about the substrate, the flow field confinement discussed in Fig. 2 and Fig. 3 phase shifts the frequency gain. The constraint on linearly stable wave numbers becomes

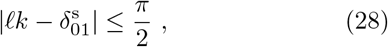

where 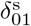 is given by Eq. (21). Fig. 5(a) illustrates that metachronal waves are linearly stable (max_*i*_Λ_*i*_(*k*) *<* 0) when the beating frequency is increased by the hydrodynamic interaction (Ω(*k*) *>* 0). The effect of flow confinement on linear stability depends on the orientation of the power stroke relative to the wave direction, as is evident by comparing the red and blue highlighted subfigures in Fig. 5(b). In particular, in the arrangement of phase oscillators modeling symplectic or antiplectic waves (blue subfigure), the flow anisotropy discussed in Fig. 2 generates a phase shift 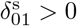. As a consequence, the number of stable wave numbers with ℓ*k >* 0 becomes larger than the number of stable wave numbers with ℓ*k <* 0. Thus, the flow confinement favors the symplectic metachronal waves. In the arrangement of phase oscillators modeling the propagation of laeoplectic or dexioplectic metachronal waves (red subfigure), the mutual flow is isotropic (Fig. 3). Thus, there is no such asymmetry between the wave directions. However, the reversal of the vertical flow with respect to the oscillator motion, discussed in Fig. 3, can induce anti-phase synchronization corresponding to 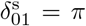. As a result, metachronal waves are stabilized at short wavelengths (around ℓ*k* = *π*) rather than at long wavelengths (around ℓ*k* = 0).

**FIG. 5.**
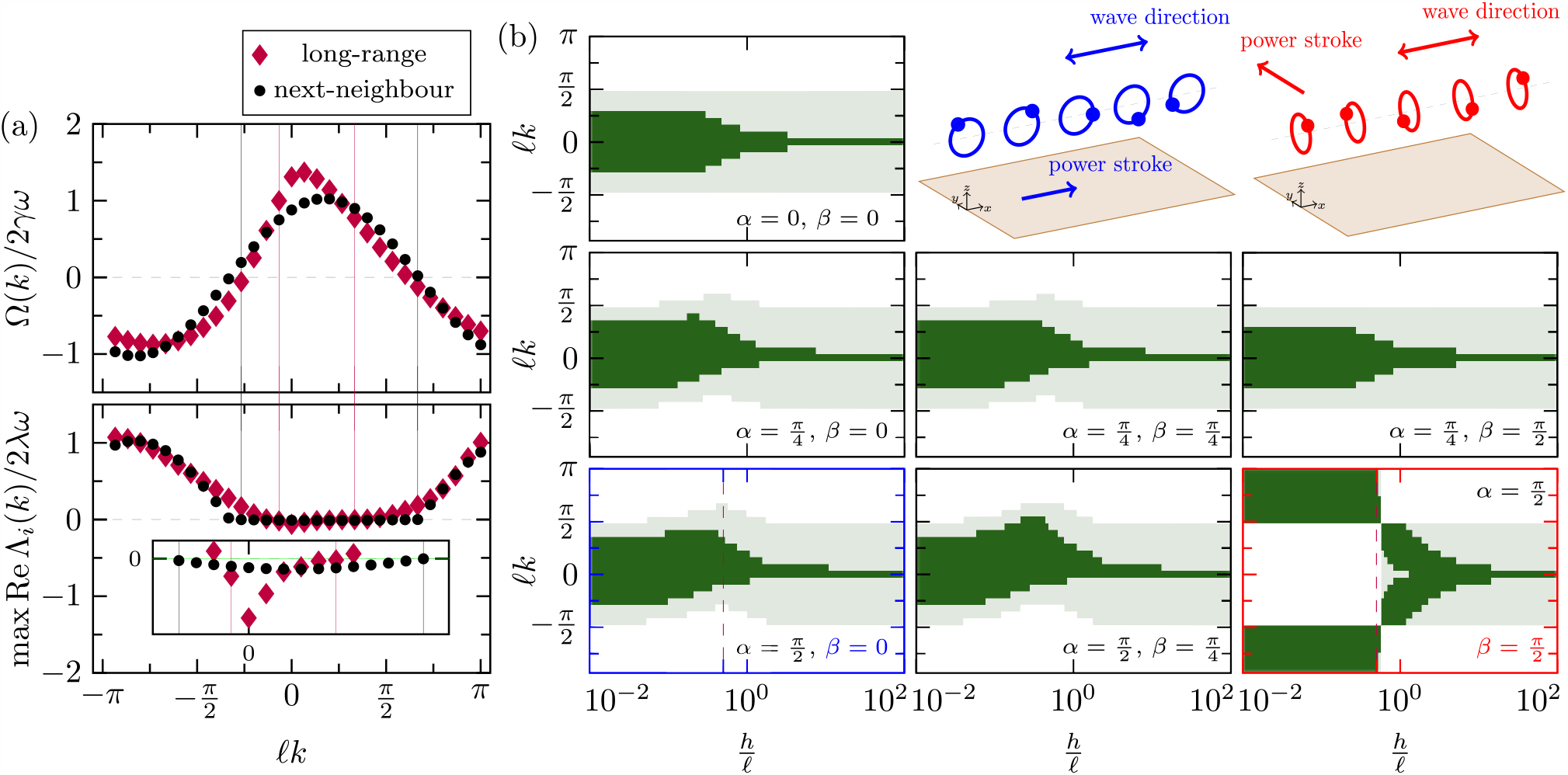
Linear stability of metachronal waves in a periodic chain of *N* = 30 phase oscillators. (a) Frequency gain Ω(*k*) and maximum linear growth rate max_*i*_ Re Λ_*i*_(*k*) for *α* = *π/*2, *β* = 0, and *h/*.ℓ = 1*/*2. Data are for next-neighbor and long-range coupling. The vertical lines indicate the linearly stable wave numbers, showing that long-range interactions reinforce the restriction to high-frequency waves that is already established by the next-neighbor interaction. (b) Linearly stable wave numbers as a function of the geometry parameters, for next-neighbor (light green) and long-range (dark green) coupling, respectively. The data for next-neighbor interactions highlights the effect of flow field confinement. As can be seen in the subfigure highlighted in blue, the linearly stable wave numbers are shifted towards positive values (symplectic waves), for a configuration as in Fig. 2, where the flow fields are anisotropic. The red highlighted subfigure shows that the long wavelengths (|ℓ*k* |*< π/*2) may be destabilized in favor of the short wavelengths (|ℓ*k*| *> π/*2), corresponding to a coupling by reversed flow fields as shown in Fig. 3. The effect of long-range coupling is seen in the collapse of the linearly stable wave numbers to the exclusively stable wave number *k* = 0, which is the maximum frequency wave number for bulk fluid hydrodynamics (*h/*.ℓ → ∞). The dashed vertical lines in (b) indicate the linearly stable wave numbers for the simulations in Fig. 6.

In the case of long-range coupling, the linear stability of metachronal waves is determined graphically via Eq. (26). Fig. 5(a) shows that an even stronger constraint on linearly stable wave numbers is imposed by long-range interactions. The effect is to increase the preference for low drag configurations, which is reflected in the fact that metachronal waves are now stabilized only near the maximum frequency wave number. How much stronger the constraint is depends on the relative weight of the long-range interactions, as can be seen by the dark green shedding of Fig. 5(b): In the *h/*ℓ → ∞ limit, the hydrodynamic interaction decays slowly with distance (∝1*/d*). Accordingly, the higher harmonics in the correction term in Eq. (26) have a significant weight, strongly restricting collective motion to low drag configurations. In fact, only the maximum frequency wave number *fk* = 0 is linearly stable. In the near-wall limit *h/*ℓ → 0, the hydrodynamic interaction is screened and the spatial decay is much faster. Consequently, the higher harmonics are less significant and multiple high-frequency wave numbers are linearly stable. In particular, in an arrangement as in Fig. 2 the coupling decays with distance ∝ 1*/d*^3^. In an arrangement like in Fig. 3 the coupling is through reversed flow fields. This leads to an even faster decay ∝ 1*/d*^5^. In this situation, long-range effects are not essential.

## IV. EMERGENCE OF METACHRONAL WAVES IN PERIODIC CHAINS

We numerically investigate briefly the spontaneous formation of metachronal waves in phase oscillator chains with periodic boundary conditions. We only consider the two basic arrangements corresponding to either symplectic (antiplectic) or laeoplectic (dexioplectic) metachronal waves. The associated linearly stable wave numbers are indicated by the dashed lines in Fig. 5(b).

To simulate extensive time series, we focus on the slow hydrodynamic interaction and remove the fast oscillations by transforming the phases as Ψ = *ϕ* − *ωt*. The dynamics of the slowly varying phases are written as

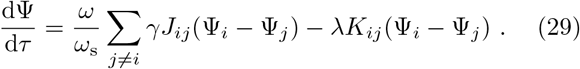

Here, we have normalized the dynamics by scaling time as *τ* = *ω*_s_*t*, where 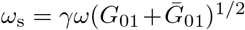. The fast phase variables are given by *ϕ*(*τ*) = Ψ(*τ*) + *τω/ω*_s_.

We obtain the data shown in the following by numerically integrating Eq. (29), by the classical Runge-Kutta method with the time step Δ*τ* = 0.01.

Fig. 6(a) shows data from simulations with 8 × 10^3^ random initial conditions, of which each converged to a metachronal wave with *ϕ*_*i*_ − *ϕ*_*i*+1_ = ℓ*k*. To quantify the emergence of metachronal waves, we calculate the probability *P* (*k*) of the wave number *k*. As expected from the linear stability analysis, the long wavelengths (|ℓ*k*| *< π/*2) emerge in the symplectic arrangement, but the short wavelengths (|ℓ*k*| *> π/*2) emerge in the laeoplectic arrangement. Note that the asymmetry between symplectic (ℓ*k >* 0) and antiplectic (ℓ*k <* 0) wave numbers – established by linear stability –, is amplified by a much larger probability of the emergence of symplectic wave numbers. The phases *ϕ*_*i*_(*t*) of typical emergent metachronal waves, over three beat cycles, are shown in Fig. 6(c) and Fig. 6(b), respectively.

**FIG. 6.**
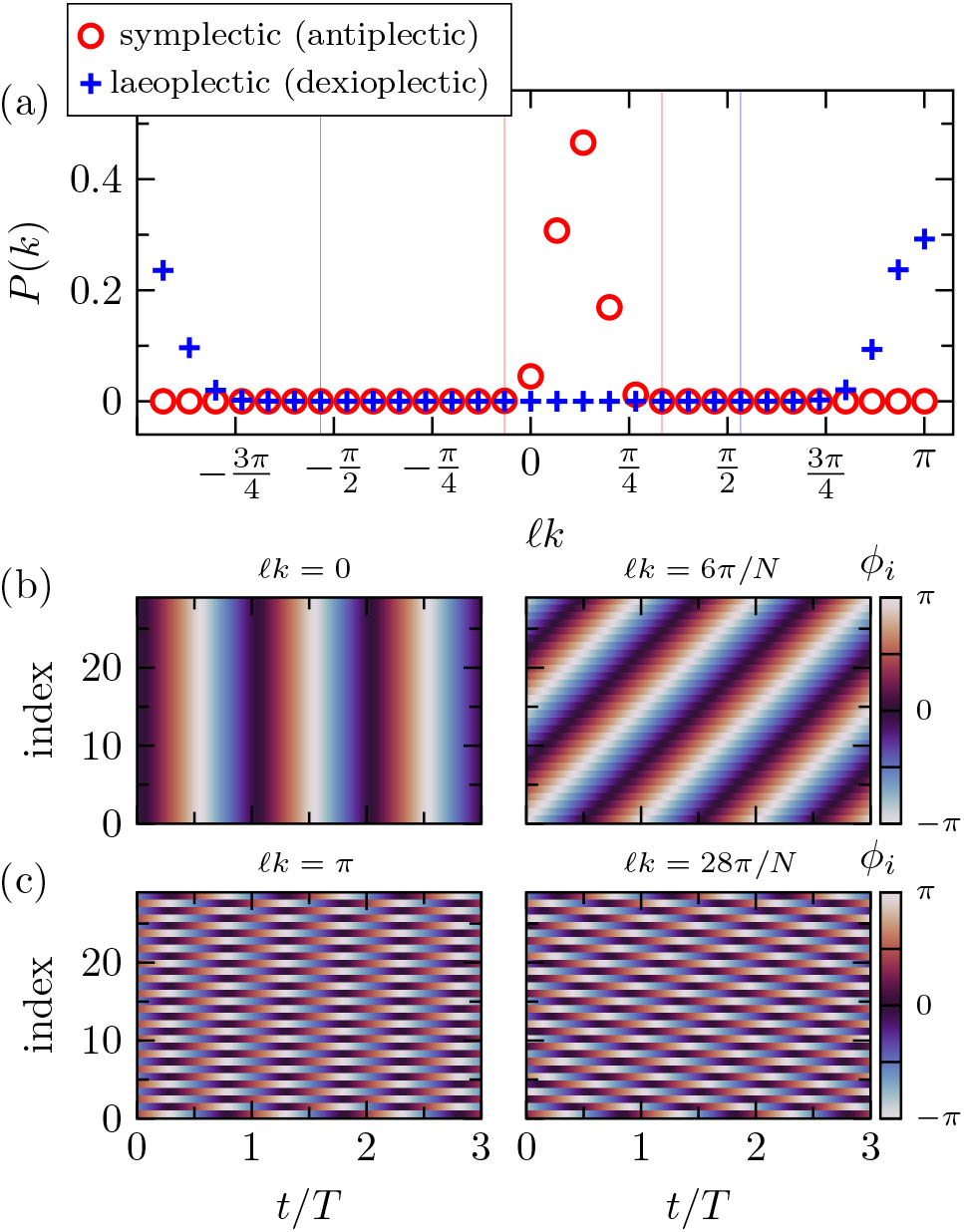
Formation of metachronal waves in periodic phase oscillator chains, oriented as in Fig. 2 (symplectic) and Fig. 3 (laeoplectic), respectively. (a) Probability *P* (*k*) of the wave number *k*. The vertical lines denote the linear stability limit according to Eq. (26). The data are from simulations with 8 *×* 10^3^ random initial conditions, each of which converged to a metachronal wave. (b)-(c) Typical phases *ϕ*_*i*_(*t*) at the end of the simulation, for the symplectic (b) and laeoplectic (c) arrangements. Parameters are *N* = 30, *h/*.ℓ = 1*/*2, *t*_e_*ω* = 1, and *γ* = 10^*−*2^. The simulation time is *τ*_sim_ = 10^3^.

## V. EMERGENCE OF METACHRONAL WAVES IN OPEN CHAINS

In many biological systems, the arrangement of cilia is not periodically closed. We investigate the emergence of metachronal waves in open phase oscillator chains by simulations of Eq. (29). We consider an arrangement of the phase oscillators as in Fig. 2, corresponding to symplectic (antiplectic) wave propagation. We choose the model parameters according to Ref. [15], where the phase-amplitude model is used as a model of metachronal waves in Volvox.

Fig. 7(a) shows the transient phases *ϕ*_*i*_(*t*), during the initial formation of a symplectic metachronal wave. As indicated by the associated transient phase shifts *ϕ*_*i*_(*t*) − *ϕ*_*i*+1_(*t*) [Fig. 7(b)], the metachronal wave is rapidly created by merging of locally phase-locked spots that are spontaneously formed by the hydrodynamic interaction. In the open chain, however, the phase oscillators at the ends interact with fewer neighbors. This prevents the system to form a collective state with uniform phase shifts *ϕ*_*i*_ − *ϕ*_*i*+1_ = *fk*, corresponding to metachronal waves in periodic chains. Nevertheless the hydrodynamic interaction leads to the formation of a metachronal wave with non-uniform steady state phase shifts, that causes frequency-locking, as shown in Fig. 7(c) and Fig. 7(e), respectively. In simulations with 10^3^ random initial conditions, we always obtained the same particular metachronal wave.

**FIG. 7.**
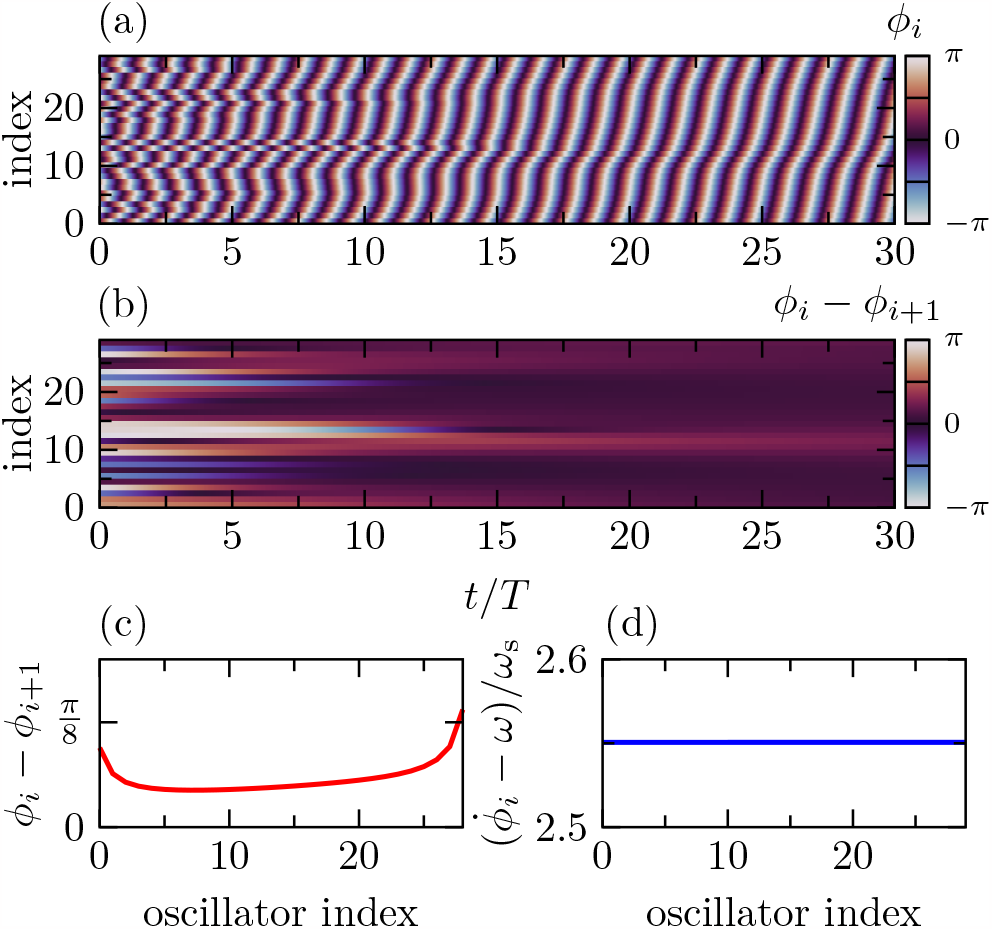
Emergence of a metachronal wave from random initial data, in an open phase oscillator chain, oriented as in Fig. 2. (a) Transient phases *ϕ*_*i*_(*t*). (b) Transient phase shifts *ϕ*_*i*_(*t*) *− ϕ*_*i*+1_(*t*). (c) Steady state phase shifts *ϕ*_*i*_ *− ϕ*_*i*+1_ after 1300 beat cycles. (d) Instantaneous frequencies 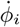 corresponding to (c). The average phase shift ⟨*ϕ*_*i*_ *− ϕ*_*i*+1_⟩_*i*_ ≈ 0.18 indicates a symplectic metachronal wave. Each random initial condition we simulated reached the same steady state. Data is generated with the parameters *h/*.*ℓ* = 1*/*2, *N* = 30, *γ* = 3*/*8 *×* 10^*−*2^, and *t*_e_*ω* = 10.

The selection of a particular metachronal wave highlights the importance of boundary conditions. To investigate how this wave selection might be influenced by the elasticity of the orbits, we continue the state shown in Fig. 7 in the elasticity range *t*_e_*ω* ∈ [10^*−*1^, 10]. Fig. 8(a)-(b) show the resulting frequency-locked solutions corresponding to either symplectic (positive phase shifts) or antiplectic (negative phase shifts) metachronal waves. By the average phase shift ⟨*ϕ*_*i*_ − *ϕ*_*i*+1_⟩ _*i*_ of the frequency-locked solutions we illustrate the elasticity dependence of the selected metachronal wave: Large elasticity corresponds to symplectic metachronal waves. Small elasticity corresponds to antiplectic metachronal waves. The transition between symplectic and antiplectic metachronal waves is characterized by an intermediate region of bistability, connecting the symplectic and antiplectic solutions by a hysteresis loop [Fig. 8(c)]. Below a critical elasticity *t*_e_*ω* ≈ 0.4, frequency-locked solutions are not observed. Thus, the instantaneous average phase shift becomes meaningless, as indicated by the irregular behavior for *t*_e_*ω <* 0.4. The phases *ϕ*_*i*_(*t*) of symplectic and antiplectic metachronal waves are shown in Fig. 8(e).

**FIG. 8.**
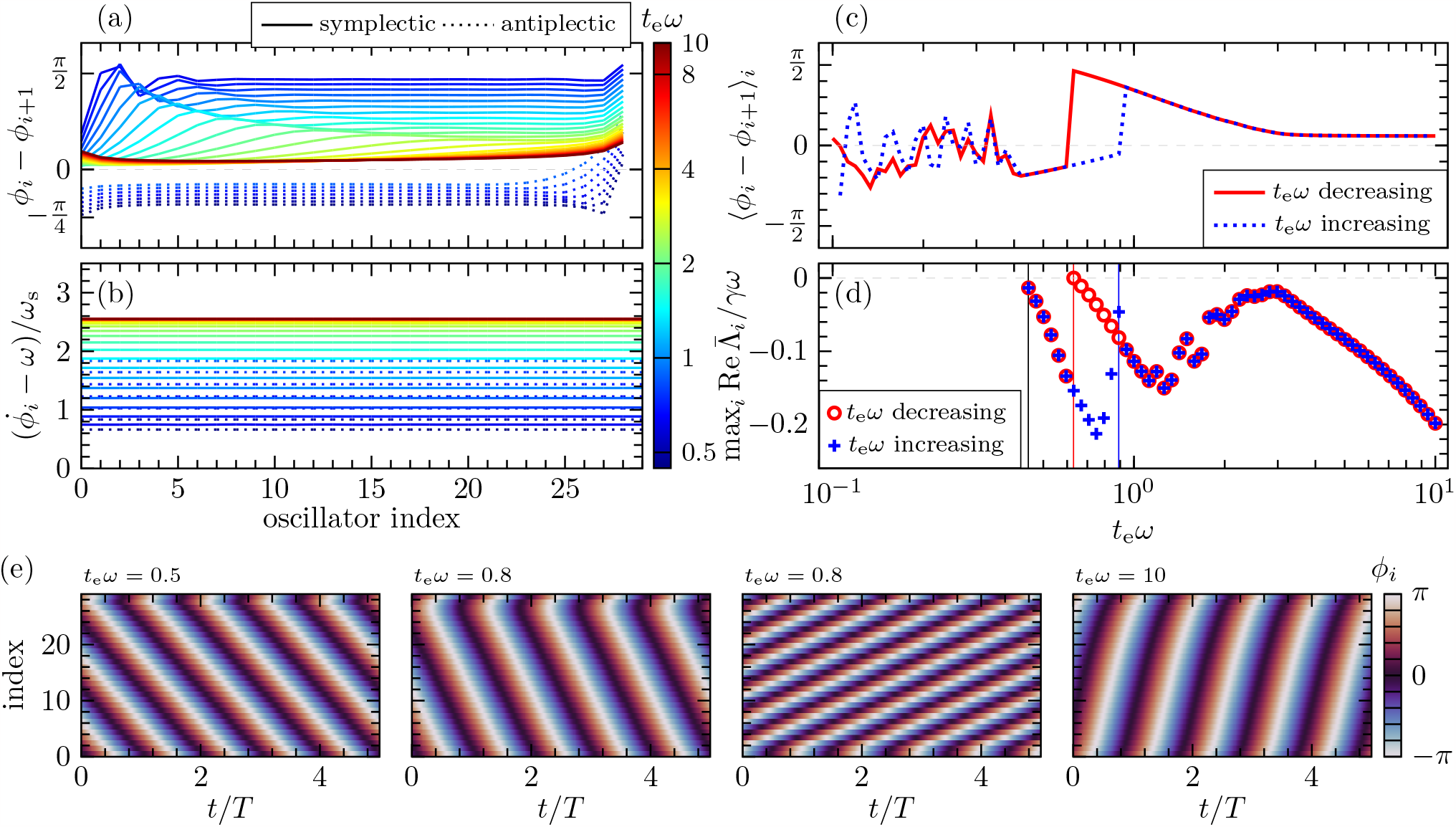
Characteristics of metachronal waves in an open chain of *N* = 30 phase oscillators, oriented as in Fig. 2 with *h/*.ℓ = 1*/*2 and *γ* = 10^*−*2^. (a) Steady state phase shifts *ϕ*_*i*_ *− ϕ*_*i*+1_ of symplectic and antiplectic metachronal waves, obtained by continuing the metachronal wave solution, shown in Fig. 7, in the range *t*_e_*ω* ∈ [10^*−*1^, 10]. (b) Corresponding frequency gain 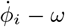 Average phase shift ⟨*ϕ*_*i*_ *− ϕ*_*i*+1_⟩_*i*_, at the end of the simulation, as a function of *t*_e_*ω*. The simulation time is *τ*_sim_ ∝ (*Nω*_s_)^2^*/*(*λω*)^2^. In the irregular regime, no steady state is reached during the simulation. (d) Maximum linear growth rate max_*i*_ Re 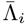 corresponding to (a). In the vicinity of the vertical lines, the growth rates are approaching a sign reversal, indicating an imminent instability. (e) Time series of the phases *ϕ*_*i*_(*t*) of metachronal waves from simulations with random initial conditions.

We obtain the linear stability of the numerically computed metachronal wave solutions similar to the periodic case: Eq. (11) is transformed into a co-rotating reference frame by 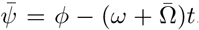, where 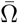 is calculated with the phase shifts in Fig. 8(a). The dynamics of 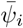 are linearized around the corresponding phase shifts. The eigenvalues 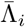 of the resulting Jacobian are calculated numerically. We discard the neutral eigenvalue – which reflects the global phase shift symmetry – when evaluating the linear stability.

Fig. 8(d) shows the maximum linear growth rate max_*i*_ Re 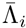, suggesting three instabilities, each occurring at an elasticity where the maximum growth rate approaches a sign reversal: (i) At the transition from symplectic to antiplectic metachronal waves (indicated by the red vertical line). (ii) At the boundary between irregular solutions and metachronal waves (black vertical line). (iii) At the transition from antiplectic to symplectic metachronal waves (blue vertical line).

While for highly elastic orbits the symplectic wave forms directly (Fig. 7), the formation process for intermediate elasticity is characterized by a competition between symplectic and antiplectic metachronal waves. We illustrate this competition in Fig. 9: The phase oscillators at the ends of the chain interact with fewer local neighbors.

**FIG. 9.**
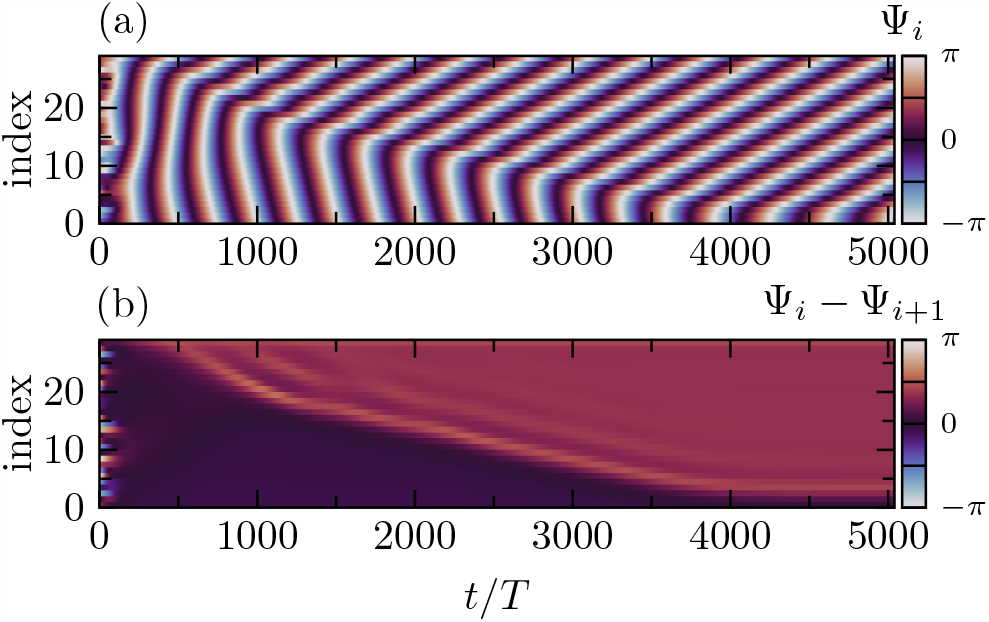
Competing symplectic and antiplectic metachronal waves in an open chain of *N* = 30 phase oscillators, oriented as in Fig. 2, with *h/*.ℓ = 1*/*2, *t*_e_*ω* = 1, and *γ* = 3*/*8 *×* 10^*−*2^. Time series of the slow phases Ψ_*i*_(*t*). (b) Phase shifts Ψ_*i*_(*t*) *−* Ψ_*i*+1_(*t*) as a function of time.

This causes a higher frequency gain of the phase oscillators in the bulk of the chain. Therefore, the phase oscillators at the ends initially lag behind their neighbors in phase. As a result, an antiplectic metachronal wave forms first in the downstream region of the chain, while the phase oscillators upstream adjust their phases to a symplectic metachronal wave. After about a thousand beats, the two metachronal waves fill the entire phase oscillator chain and intersect at a synchronized interface. This pattern of two oppositely directed metachronal waves is sometimes called a chevron pattern. For bulk fluid hydrodynamics, the chevron is a stable collective configuration [34]. In our model, however, the phase oscillators interact through anisotropic near-wall flows that implement a directional coupling. The two-wave state is no longer stable, and the symplectic and antiplectic metachronal waves compete to push each other out of the phase oscillator chain, approaching a single-wave frequency-locked solution on a time scale of several thousand beat cycles. This dynamic behavior is consistent with a discontinuous transition between an antiplectic wave and a symplectic wave [Fig. 8(c)].

## VI. SUMMARY AND DISCUSSION

In this paper, we investigate hydrodynamic synchronization between cilia which are modeled by phase oscillators whose elastic circular orbits are inclined about a no-slip wall. The inclination of the orbit breaks the symmetry of the circular motion and establishes a preferred direction of flow, thus modeling the power stroke of the cilia. The power stroke direction in turn leads to an anisotropy in the hydrodynamics, breaking the reflection symmetry between upstream and downstream neighbors and reversing the coupling with respect to mutual motion for oscillators oriented perpendicular to the power stroke. The anisotropic hydrodynamics results in directional coupling, which has profound consequences for the coordination of metachronal waves: In periodic phase oscillator chains, metachronal waves propagating in the direction of the power stroke are linearly stable at long wavelengths. The perpendicular wave directions, however, may be stable at short wavelengths. Furthermore, the interplay between flow confinement and orbit compliance robustly coordinates the phase oscillators in metachronal waves independent of the boundary and initial conditions. In open chains of phase oscillators, we observe a phenomenon which is novel in the context of hydrodynamic synchronization of cilia: Bistability of symplectic and antiplectic metachronal waves leads to hysteresis. Adjusting the elasticity allows to switch between the wave directions.

Hydrodynamic interactions between inclined, elastically bound, harmonically oscillating microspheres have previously been studied numerically in terms of coupled phase-amplitude oscillators [14, 15]. Our model exhibits essentially identical coupling characteristics between pairs of oscillators (see App. C), particularly with respect to the spatial decay of the coupling. The models should therefore predict similar large scale coordination. Yet, in simulations with the phase-amplitude model over about a thousand beat cycles, bistability of metachronal wave directions is not reported [14, 15]. In our simulations, the time scale of competition between wave directions is rather slow, requiring several thousand beat cycles.

The geometrical understanding of the hydrodynamic coupling, illustrated in Fig. 2 and Fig. 3, may have relevance beyond coupled phase oscillators. The discussed effects of flow confinement solely depend on the presence of a boundary that confines the flow. Thus, slender filaments undergoing complex motion are expected to produce a flow that exhibits similar flow confinement. Indeed, long-wavelength symplectic metachronal waves and short-wavelength laeoplectic metachronal waves are observed in an in vitro model of ciliated bronchial epithelium [19]. Flow confinement could explain these observations. However, to further link the interplay of flow confinement and elastic compliance with metachronal co-ordination in biology, it is necessary to extend our analysis to carpets of phase oscillators.

In experiments with laser-driven colloids [57] as well as in numerical investigations [58] an instability of metachronal waves with finite wavelength occurs for interactions far from the no-slip wall. The linear stability of metachronal waves in our model suggests that this instability is caused by a constraint for low-drag collective configurations induced by long-range interactions, which ultimately stabilizes only the highest frequency wave mode, corresponding to an infinite wavelength in the bulk fluid. However, conclusions about long-range hydrodynamic effects derived from models based on the steady Stokes equations must be treated with caution, since these equations break down at large length scales (compared to the length scale of vorticity diffusion 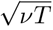) [59]. In fact, recent experiments suggest that inertial effects in the fluid cannot be neglected at the microscale [60, 61]. Furthermore, theory suggests that the leading order contribution to the hydrodynamic synchronization strength, induced by time dependent effects in the fluid, scales with distance fundamentally different then expected from steady Stokes flow [39]. To judge possible deviations from the steady Stokes flow coupling in systems extending beyond a length scale 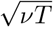, a theory is needed that takes into account the time dependence of the fluid.

Elastic compliance and variable forcing can similarly contribute to hydrodynamic synchronization [42], and their interrelationships can be used to control synchronization between oscillator pairs [41]. Regarding hydrodynamic large-scale coordination, we note that, as shown in Sec. III, the dispersion of metachronal waves in the elastic compliance framework is dominated by first harmonics. In contrast, the dispersion of metachronal waves coordinated by variable driving forces is dominated by the second harmonics [46]. This suggests that metachronal waves are controllable by relative adjustments of the elastic parameter and the coefficients of variable forcing.

## ACKNOWLEDGMENTS

We acknowledge funding by the Deutsche Forschungsgemeinschaft trough the Collaborative Research Center 910.

## Appendix A: Phase oscillator dynamics

Here, we derive the coupled phase oscillator equations at the time scale of synchronization to the leading order in the hydrodynamic interaction.

The position of the model cilia reads as

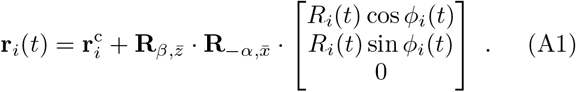

where 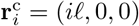 with *i* ∈ (0, 1, …, *N* − 1). The circular orbits are locally rotated by the matrices

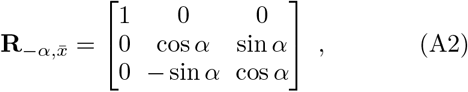

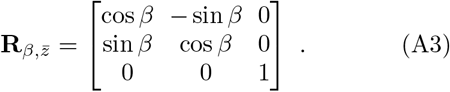

The tangential and radial unit vectors are given by

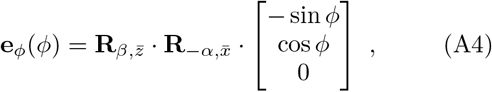

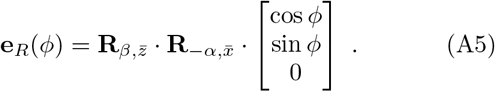

The dynamic equations of the phase and the radius can be written as

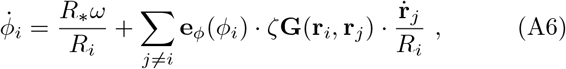

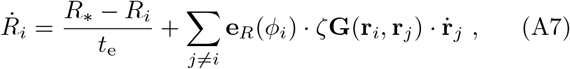

The Greens function **G** of the steady Stokes equation, subject to a no-slip condition on the substrate, is known as the Blake tensor [55]. The components of the Blake tensor (*μ, ν* ∈ {*x, y, z*}) can be written as

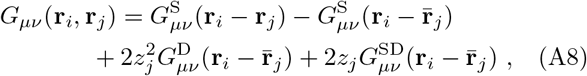

with 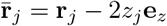 and

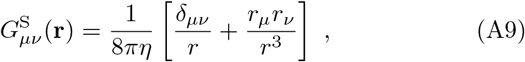

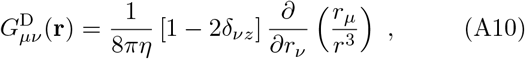

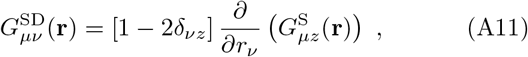

where *r* = |**r**| and *d*_*μν*_ denotes the Kronecker delta [62].

To simplify the Blake tensor we neglect the finite size of the orbits and assume the scaling separations *R* ≪ ℓ and *R* ≪ *h*. The distance vectors read as

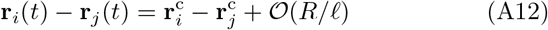

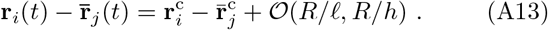

Furthermore, we decompose the Blake tensor in symmetric and anti-symmetric parts as

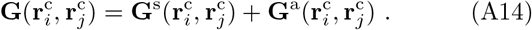

As basis for the decomposition we use the center to center unit vectors 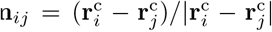 and 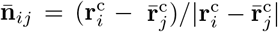. Those are given by

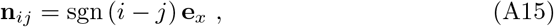

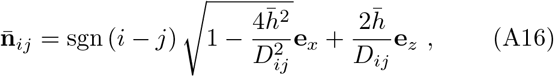

where 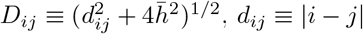 and 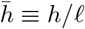.

In order to generalize our model to carpets of cilia this basis vectors must be modified.

The symmetric and anti-symmetric parts of the Blake tensor can be written as

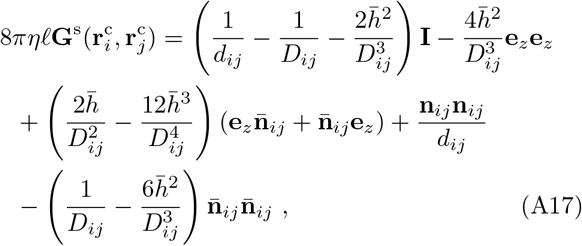

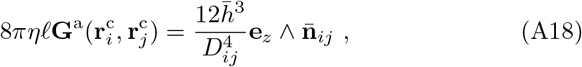

where ***ab*** = *a*_*μ*_*b*_*ν*_ and ***a*** ∧ ***b*** = ***ab*** − ***ba***.

With *ζ* = 6*πηa* and the factor 8*πηf* in Eqs. (A17)-(A18), we see that the interaction terms in Eqs. (A6)-(A7) scale as *ζ***G** = 𝒪 (3*a/*4ℓ). We only consider weak hydrodynamic interactions with *γ* ≡ 3*a/*4ℓ ≪ 1.

Starting form an unperturbed motion 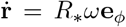 in Eq. (A7), we estimate the scale of the hydrodynamically induced radial velocity and the radial displacement as 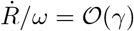 and *R/R*_*∗*_ = 1+O(*λ*), where *λ* = *γt*_e_*ω* ≪ 1. The resulting changes in the velocity 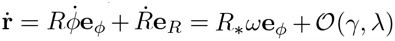 induce interaction terms of the order of *γ*^2^ and *λγ* in Eqs. (A6)-(A7).

With Eq. (A14) and keeping only the leading order in *λ, γ*, Eqs. (A6)-(A7) reduce to

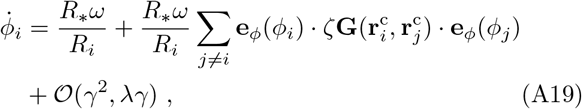

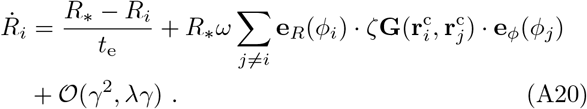

Using Eqs. (A4)-(A5) and Eqs. (A14)-(A18) we calculate the pair wise hydrodynamic coupling functions as

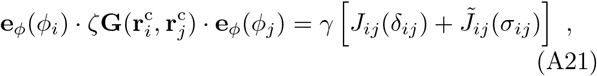

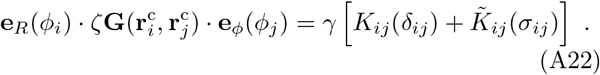

where *d*_*ij*_ ≡ *ϕ*_*i*_ − *ϕ*_*j*_, *σ*_*ij*_ ≡ *ϕ*_*i*_ + *ϕ*_*j*_ and

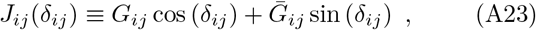

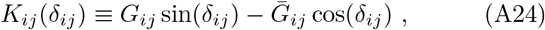

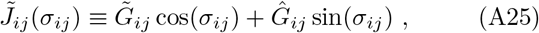

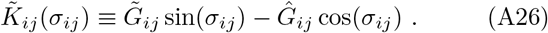

The geometry factors read

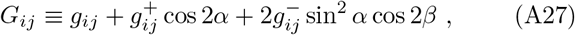

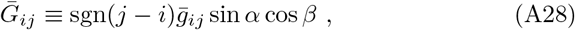

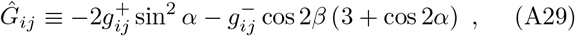

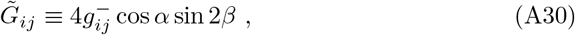

with dimensionless distance functions

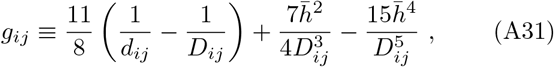

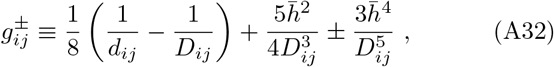

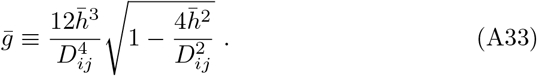

The phase difference changes at the time scale of synchronization 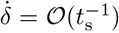, but the sum of phases changes at the order of the intrinsic frequency 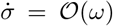 [34].

Thus, Eqs. (A21)-(A22) show that the hydrodynamic interaction separates into slowly and rapidly varying parts. We are only interested in the synchronization dynamics and remove the rapid oscillations by period averaging as

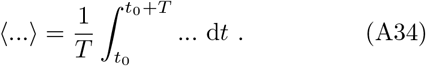

We assume that the synchronization time is considerably larger then the period (*t*_s_ » *T*) and take *δ* as constant over one period. We moreover ignore the higher order terms in *λ, γ* and set *σ* = 2*ωt* + 𝒪 (*γ, λ*) in the averaging of the coupling functions. The average radial dynamics are given by

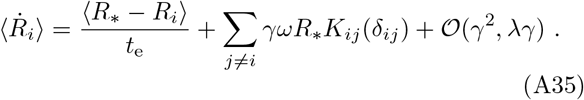

The first term on the right hand side of Eq. (A35) generates a relaxation to an average radial displacement at a time scale *t*_e_. The second term represents the hydrodynamic coupling, which slowly changes at the time scale of synchronization *t*_s_.

To the leading order in the radial displacements the average phase velocity reads as

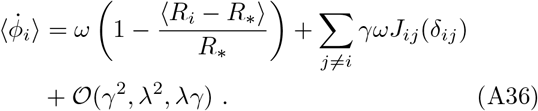

To arrive at the phase oscillator equation we exploit the scaling separation *t*_e_ ≪ *t*_s_ and set 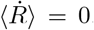. Then, by Eq. (A35), the phase dynamics at the synchronization time scale can be written as

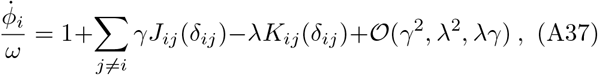

where we dropped the averaging indication.

A comparison between the full system of phase and radial dynamics Eqs. (A19)-(A20) and our phase oscillator approximation is shown in Fig. 10. If *t*_e_*/t*_s_ ≈ 1 the phase difference becomes modulated at a fast time scale, leading to optimal synchronization at finite elasticity [56]. In this case our approximation breaks down.

**FIG. 10.**
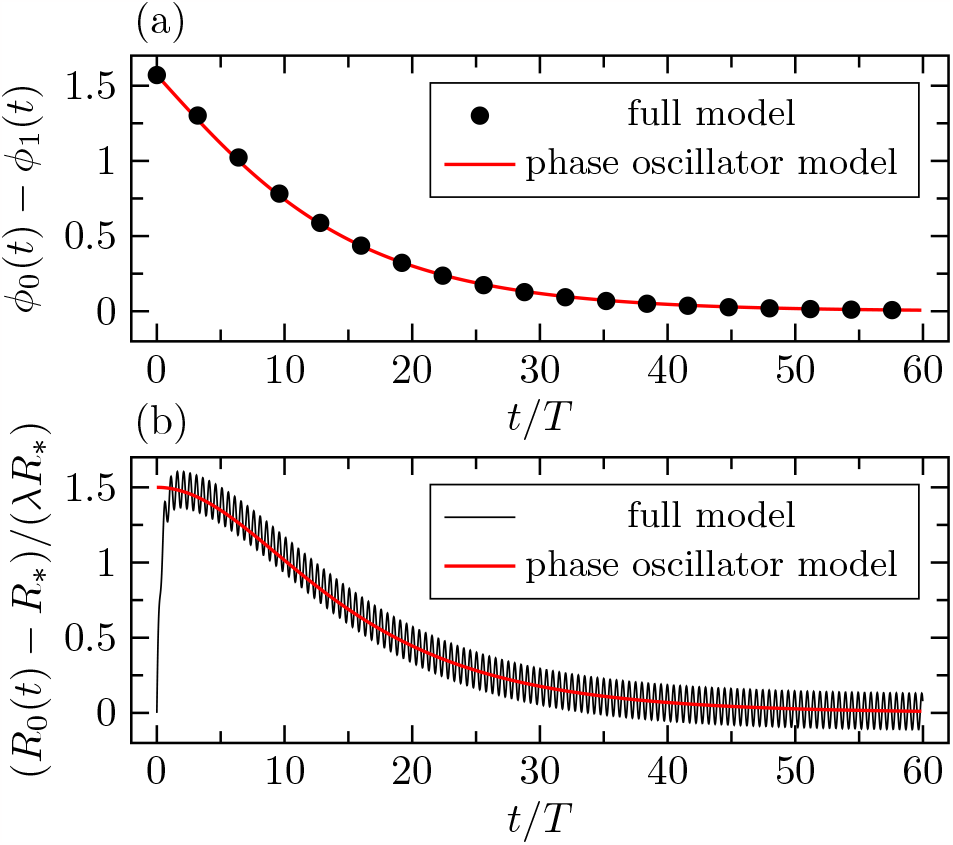
We numerically solve Eqs. (A19)-(A20), for a pair of phase oscillators, in the bulk fluid (*h/*.ℓ → ∞), by the classical Runge-Kutta method with numerical step size Δ*t* = 0.01. (a) Phase difference *ϕ*_0_(*t*) *− ϕ*_1_(*t*) according to the full model and according to our phase oscillator model [Eq. (C6)]. (b) Numerically calculated radial displacement *R*_0_(*t*) *− R*_*∗*_ and average radial displacement according to Eq. (A35) (with 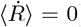 and Eq. (C6). Initial data are *R*_0_(0) = *R*_1_(0) = *R*_*∗*_ and *ϕ*_0_(0) *− ϕ*_1_(0) = *π/*2. Parameters are *α* = *β* = 0, *γ* = 1*/*4 *×* 10^*−*2^, and *t*_e_*ω* = 2.

## Appendix B: Linear growth rate of perturbations of metachronal waves in periodic chains

The Jacobi matrix of metachronal waves in periodic chains of *N* phase oscillators is given by

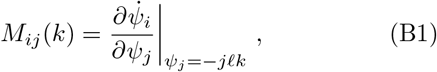

with |ℓ*k*| = 2*πK/N*, where *K* is an integer and

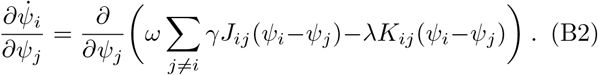

We impose a cutoff radius of half the system size.

Due to the periodic boundary conditions and the translation invariance of the phase shifts of metachronal waves the Jacobi matrix is a circulant matrix; its successive rows are given by cyclic right shifts of its first row. Thus, the matrix is fully specified by the components *M*_0*j*_, say.

We begin by calculating *M*_00_ as

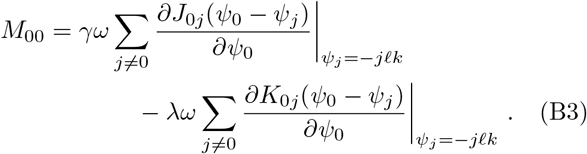

The interaction ranges symmetrically to either side of the periodic phase oscillator chain. We split the summation in two sums. One for each direction. This leads to

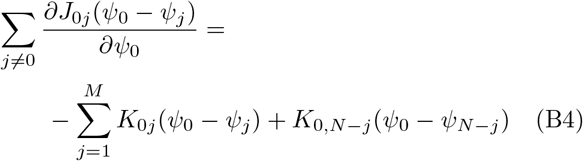

and

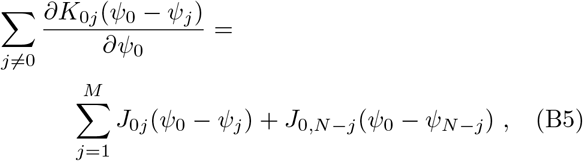

where *M* = (*N* − 1)*/*2 for odd *N* and *M* = *N/*2 − 1 for even *N* .

The *j*th neighbor phase difference is *ψ*_*i*_ −*ψ*_*i*+*j*_ = *jℓk* in one direction and *ψ*_*i*_−*ψ*_*i−j*_ = −*jℓk* in the other direction. Thus, in the wave state we can identify *ψ*_0_ −*ψ*_*j*_ = − *jℓk* and *ψ*_0_ − *ψ*_*N−j*_ = −*jℓk*. At the level of the hydrodynamic coupling functions we put *J*_0*j*_(*ψ*_0_ − *ψ*_*j*_) = *J*_0*j*_(*jℓk*) in one direction and *J*_0,*N−j*_(*ψ*_0_ − *ψ*_*N−j*_) = *J*_*j*0_(− *jℓk*) in the other direction. The same holds for the coupling function *K*_*ij*_. With this we arrive at

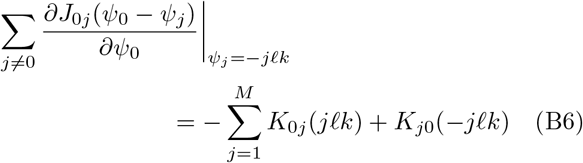

and

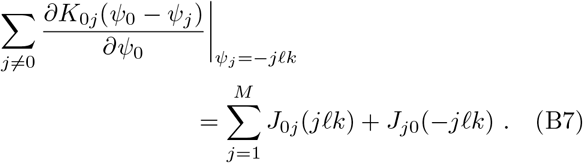

Using Eqs. (B6)-(B7) and Eqs. (19)-(20) in Eq. (B3) we obtain

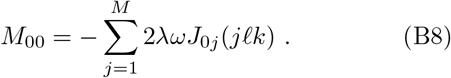

The *j* ≠ 0 components of 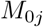 are given by

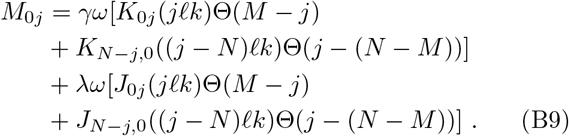

where Θ(*x*) is the Heaviside step function. The eigenvalues Λ_*n*_ are written as

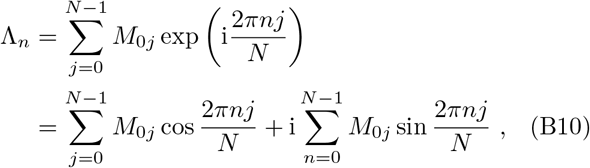

where 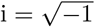 [63]. The real part reads as

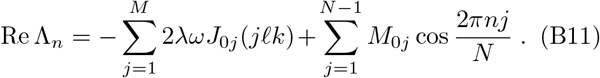

After carrying out the summation over 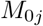 and rearranging indices, the second term in Eq. (B11) reads

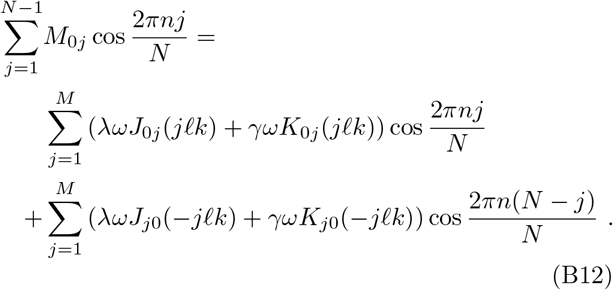

Using Eqs. (19)-(20) in Eq. (B12), we obtain the linear growth rates from Eq. (B11) as

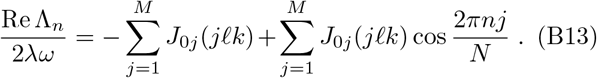

## Appendix C: Pair synchronization

Here, we compare our phase oscillator model with the more general phase-amplitude model. In Ref. [15], it is shown that the phase difference Δ = ϕ_1_ − ϕ_0_ in the phase-amplitude model obeys the evolution equation 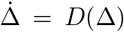, where the numerically computed coupling function is given to a good approximation by *D*(Δ) = − *D*_0_ sin(Δ− Δ_s_), with amplitude *D*_0_ and phase shift Δ_s_. Similarly the sum of phases Σ = ϕ_0_ + ϕ_1_ evolves according to 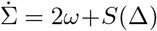, where *S*(Δ) = *S*_0_ cos(Δ− Δ_s_). It is shown that the phase shift Δ_s_ and the amplitudes *D*_0_, *S*_0_ depend in a particular way on the geometrical configuration, leading to in-phase motion, anti-phase motion, and phase-shifted steady states, depending on the orientation of the oscillators relative to the direction of the power stroke.

In our model the dynamic equations of the phase difference *δ* = *ϕ*_0_ − *ϕ*_1_ and the sum of phases *σ* = *ϕ*_0_ + *ϕ*_1_ are given by

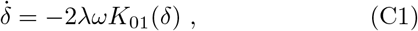

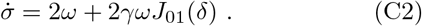

By Eq. (12) and Eq. (18) it is clear that the functional form of the dynamics above coincide with the evolution equations of the phase-amplitude model. In particular we can write 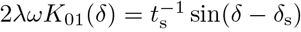 sin(*d* − *d*_s_), where the steady state phase shift *δ*_s_ and the synchronization time scale *t*_s_ are given by

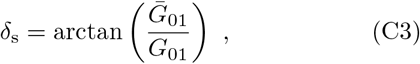

and

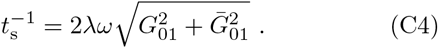

Similarly we write 2*ωγJ*_01_(*d*) = 2*ω*_s_ cos(*d* −*d*_s_), where the steady state frequency gain 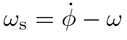 reads as

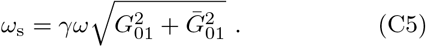

Furthermore, we solve Eq. (C1) explicitly for a function of time *δ*(*t*) as

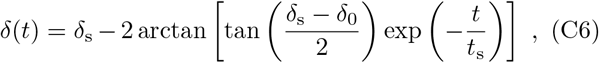

where *d*_0_ is the initial phase difference.

The general properties of the hydrodynamic synchronization between pairs of phase oscillators are shown in

Fig. 11. We explicitly reproduce the data of the phaseamplitude model shown in Fig.5 of Ref. [15]. A direct comparison between Fig. 11 and Fig.5 of Ref. [15] shows that the coupling of the phase-amplitude model agrees remarkably well with the coupling of our phase oscillator model.

**FIG. 11.**
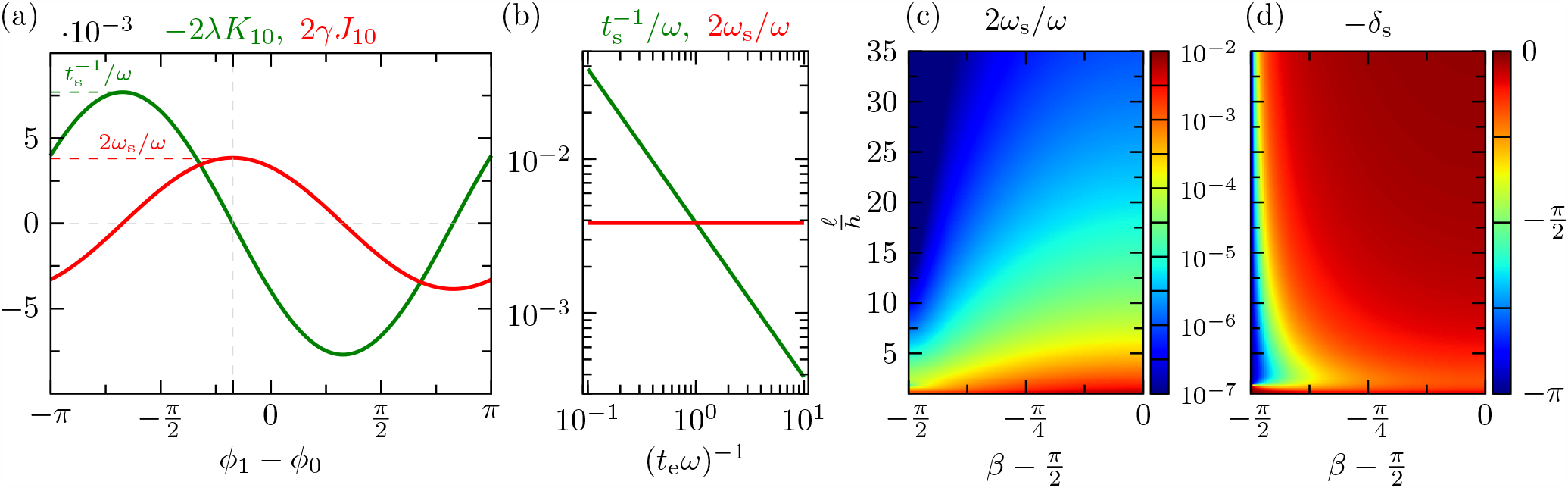
Synchronization between pairs of phase oscillators. (a) Cycle-averaged hydrodynamic coupling functions as a function of phase difference. (b) Synchronization time *t*_s_ and frequency gain *ω*_s_ as a function of the stiffness of the orbits. (c) Steady state phase difference *−d*_s_ as a function of the geometric configuration. (d) Frequency gain *ω*_s_ as a function of the geometric configuration. The data shown corresponds to the data in Fig. 5 of Ref. [15]. In order to facilitate the comparison we shifted the orientation parameter *β* and show data for *−d*_s_. Unless otherwise stated the data are generated with the parameters *α* = *π/*2, *β* = 0, . ℓ*/h* = 2, *γ* = 3*/*8 *×* 10^*−*2^, and *t*_e_*ω* = 2.

